# Ensembl Core Software Resources: storage and programmatic access for DNA sequence and genome annotation

**DOI:** 10.1101/087239

**Authors:** Magali Ruffier, Andreas Kähäri, Monika Komorowska, Stephen Keenan, Matthew R. Laird, Ian Longden, Glenn Proctor, Searle Steve, Daniel Staines, Kieron Taylor, Alessandro Vullo, Andrew Yates, Daniel Zerbino, Paul Flicek

## Abstract

The Ensembl software resources are a stable infrastructure to store, access and manipulate genome assemblies and their functional annotations. The Ensembl “Core” database and Application Programming Interface (API) was our first major piece of software infrastructure and remains at the centre of all of our genome resources. Since its initial design more than fifteen years ago, the number of publicly available genomic, transcriptomic and proteomic datasets has grown enormously, accelerated by continuous advances in DNA sequencing technology. Initially intended to provide annotation for the reference human genome, we have extended our framework to support the genomes of all species as well as richer assembly models. Cross-referenced links to other informatics resources facilitate searching our database with a variety of popular identifiers such as UniProt and RefSeq. Our comprehensive and robust framework storing a large diversity of genome annotations in one location serves as a platform for other groups to generate and maintain their own tailored annotation. Our databases and APIs are publicly available and all of our source code is released with a permissive Apache v2.0 licence at http://github.com/Ensembl.

## Introduction

The Ensembl database infrastructure was originally designed to support the storage and distribution of the reference assembly produced by the Human Genome Project (HGP) (1). Today, for any species or clade of interest, an Ensembl Core MySQL relational database can store the assembly structure, genomic sequence and genome annotations. Other Ensembl databases specialise in variation and phenotype data (2); whole genome alignments and other comparative genomics information (3); and epigenomic data and regulatory annotation (4).

Application Programming Interfaces (APIs) implemented in Perl provide primary access to these databases. The APIs offer simple functions to manage database connections, formulate queries and provide an object relational mapping (ORM) layer (i.e. they convert genomic annotations stored in a database into Perl objects). Key biological concepts such as genes, transcripts and exons are thus modelled as specialised objects, with links to their components and other related elements. In contrast, simpler features such as CpG islands can be stored as generic region objects located on the genome. Thus, the Ensembl Core database and API is the foundation for all Ensembl data resources (3–7) as well as our web browser, http://www.ensembl.org.

The number, complexity and diversity of available genome assemblies has been in constant growth since the early days of Ensembl. For example, our previous description of the Ensembl Core software libraries included a schema to represent genome assemblies resulting from the clone-by-clone based sequencing strategy used in the HGP (8), which was rendered intractable by whole genome shotgun based assembly methods. Starting with the human genome, sequencing projects focused on known haplotypes of the major histocompatibility complex (MHC) (9) or global population variation (10) captured genetic diversity, while essential curation has progressively identified and fixed errors in the reference assembly without updating the entire assembly version (11). Related variation discovery and targeted loci studies have been conducted in a number of species (12–14). At the same time the genomes of many species from across the phylogenetic tree have been sequenced and released, including tens of thousands of bacterial genomes (15). In short, the number, complexity and diversity of available genome assemblies is in constant growth.

The growing number of genome sequences has been accompanied by a substantial growth in genomics data resources (16). Established resources such as UniProt (17), RefSeq (18) and the HUGO Gene Nomenclature Committee (HGNC) (19) have increased in depth and value. Their identifiers and terminology have become a critical *lingua franca* that ties together the field and makes the results of diverse studies quickly and effectively comparable.

To keep up with these developments, the Ensembl Core database and API infrastructure has been extended in many ways. It efficiently manages whole genome shotgun genome assemblies and many species within a single database. It represents modern assembly features, beyond consensus haploid sequence, including alternate sequences, pseudo-autosomal regions (PAR) and assembly corrections (patches), while maintaining a minimal data footprint. Methods to map stored gene identifiers between releases and to link them to external databases are more robust. There exists native API support for polycistronic transcripts and ontologies. We have also adopted alternative storage solutions where our MySQL database approach does not scale for particular data types such as RNA-seq read alignments (20).

We describe here our current methods for representing genome assemblies and annotation for all species. We detail the way we manage transitions of assembly coordinates and stable identifiers (i.e. Ensembl gene, transcript, exon and other ids) across releases. Improvements to the Core Perl API and our methods for cross-referencing identifiers from external databases will also be described.

## Methods

### Representing genomes

The original Ensembl Core database schema was designed for data from the draft releases of the HGP. As a consequence, the schema modelled the common units of that assembly including bacterial artificial chromosome (BAC) clones, contigs and chromosomes (8). This model made a number of assumptions: chromosomes were necessarily composed of contigs, which were necessarily composed of clone sequences. The design became obsolete in various ways: supercontigs could not be modelled; all DNA sequence had to be assigned to clones; and a database could only hold a single assembly version.

The schema was remodelled in Ensembl release 20 into six tables as shown in Figure 1. An assembly is now composed of *sequence regions*. Sequence regions are grouped by coordinate system (e.g. chromosome, contig, plasmid), an optional assembly version and whether they have DNA sequence associated with them. The *assembly* table defines a hierarchy of sequence regions: any sequence region can be decomposed into component sequence regions much like an AGP file (https://www.ncbi.nlm.nih.gov/assembly/agp/AGP_Specification/) or INSDC CON (21) record. The *seqlevel* coordinate system is at the bottom of each assembly and refers to regions with DNA sequence (usually contigs). The *toplevel* coordinate system represents all the regions that cannot be collected into larger regions. Typically, chromosomes are *toplevel* sequences, as are supercontigs, scaffolds or contigs that have not been mapped to chromosomes. Genomic annotations are stored on *toplevel* sequence regions for best query performance.

**Figure 1:**
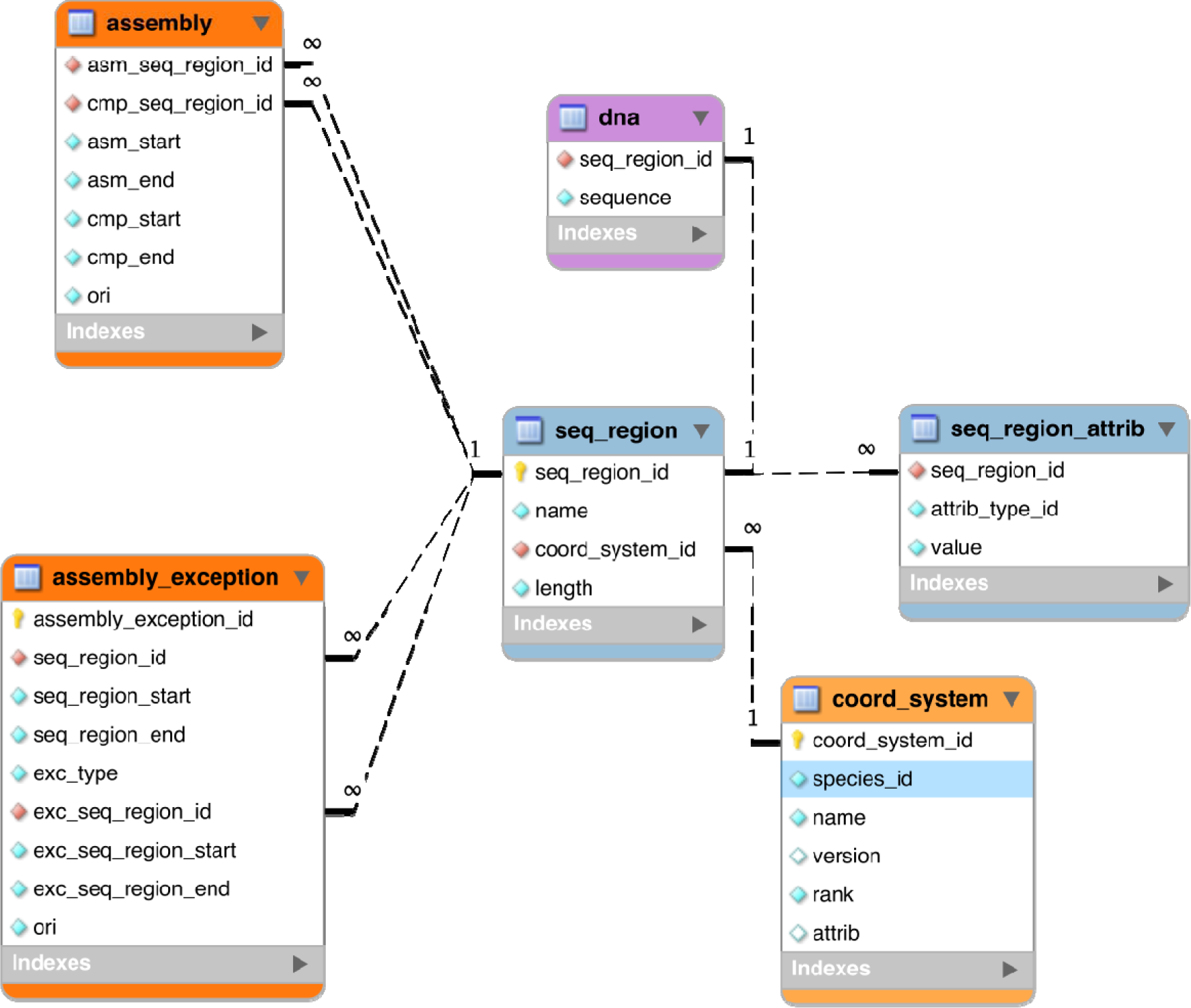
The core assembly schema.

The reference genome has been enriched with a set of ancillary sequences, which we describe using our “assembly exception” model. Assembly exceptions allow for the efficient storage of alternate loci as well as novel and patch sequences from the Genome Reference Consortium (GRC) (22) by only keeping the changes required to recreate them. Alternate loci are separate representations of regions found in the primary assembly such as the human MHC regions on chromosome 6. Patches either fix errors on the genome assembly or add novel sequence. For example, a patch on chromosome 9 of GRCh38 fixes a local mis-assembly and allows Ensembl to present the correct annotation for the ABO locus (ENSG00000175164) on a patch (KN196479.1).

Although the underlying sequence data is stored without duplication, assembly exceptions are presented as whole chromosomes where the key region (i.e. alternate haplotype, sequence patch, etc.) has been replaced by different sequence. Thus any feature coordinates on these patches are computed relative to the entire chromosome rather than to the modified region. Similarly, annotated features that are downstream of the exception may have modified coordinates when queried in the context of exception sequence.

For practical reasons, any annotation on the chromosome that is located across an assembly exception is duplicated and stored for both the primary chromosome and on the exception. In particular, multiple copies of the same gene are modelled as separate gene records, some of which may only be found on alternative assembly regions (Figure 2). The resulting multiple gene copies are called alternative alleles, or *alt alleles*, and collected together as a group of genes with a nominated reference copy. The alt alleles data structure also stores information about whether the duplicate annotation was due to sequencing error or individual variability and whether it was assigned automatically or manually.

**Figure 2:**
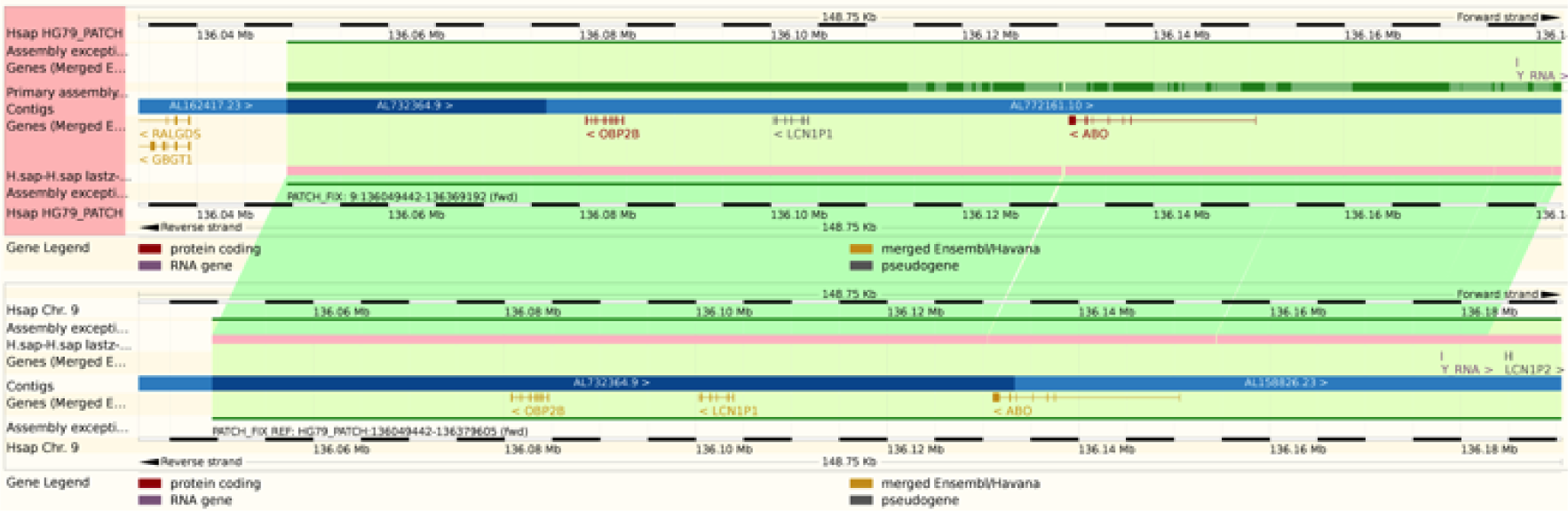
The Ensembl web browser can display the differences between a patch region and its equivalent in the primary assembly. Genes that are present in both regions are identified as alt_alleles.

Querying across an assembly exception requires searching separately the main assembly, upstream and downstream of the assembly exception location, and the assembly exception itself. When querying the flanking regions our API automatically handles these operations and discards all duplicated features.

We also use the assembly exception concept to store the pseudo-autosomal regions (PARs), which are the relatively short regions that behave as homologous autosomal regions within the otherwise highly differentiated mammalian sex chromosomes (23). For example, the human Y chromosome shares two regions (10001-2781479 and 56887903-57217415 on the GRCh38 assembly) with the X chromosome. One copy of the annotation and sequence is stored on X with equivalent regions on Y marked as assembly exceptions. When querying across these regions in Y, our API automatically projects any available genomic annotation on X to the corresponding coordinate space on the Y chromosome.

## Perl API

The Perl API is a powerful tool to store and retrieve genome annotation. The infrastructure relies on a clean separation between data representation and data access. All biological features including genes, transcripts and exons have a dedicated data object describing their attributes and logic. An adaptor class is responsible for querying the underlying databases providing a clean separation between business logic and database persistence and insulating API users from underlying schema changes. Data objects and adaptors are named after the entity they model or retrieve. For example, genes are modelled by the Gene object and are retrieved via the GeneAdaptor. In addition data objects are capable of retrieving additional records from the database via the adaptor framework allowing a number of attributes—such as all transcripts linked to a gene—to be loaded on-demand. Here we present some of the API’s characteristics.

### Genomic Features

All genomic annotations share some common attributes, such as start and end position on a sequence region, and are defined by the Feature class. More specialised features are derived from this class and include regulatory features, variants, constrained elements and sequence repeats. The base Feature class offers key operations to retrieve co-located features of a genomic region, map features between assemblies, test object equality, detect feature overlap and project features onto assembly exceptions.

The feature object hierarchy works in parallel to the adaptor hierarchy rooted by BaseFeatureAdaptor, which provides a framework for developing feature ORMs. The base class implements optimised SQL generation and automatic region expansion should a query span an assembly exception or when features are located on another coordinate system. If required, the coordinates of a retrieved feature are automatically recalculated providing locations relative to the queried region. All derived feature adaptors implement a number of stub methods that specify the tables to query, columns to retrieve and query optimisation hints.

Features link back to their sequence region via objects of class Slice, which are portions of sequence regions with a coordinate system characterised by a start, end and orientation. The Slice class also provides a number of shortcut methods to retrieve sequence (repeat masked and reverse complemented) and co-located genomic features. Slice is therefore used as the main entry point to query for annotation by genomic position and holds a number of convenience methods for multiple data set retrieval.

### Efficient Feature Searching

A major obstacle when storing genomic features in a database is how to ensure the efficient querying of randomly accessed regions. As table sizes tend towards millions of entries, naive solutions scale poorly. Genome binning schemes such as those used by the UCSC Genome Browser (24) and the BAM file format (25) partly address these problems.

We have chosen an alternative indexing strategy: all feature tables are sorted by their sequence region identifier and start coordinate, and a B-tree index is generated over the same fields. Storing related data sequentially ensures that disk seeks are minimised during regional queries. Furthermore, we compute for each table and each coordinate system the maximum feature length and store this value in the *meta_coord* table. For each genomic region query, a maximal search window for possible start positions is calculated based on the known maximum feature length and the region’s start (Figure 3) and queried against the B-tree index. This query will return all data that overlaps the queried genomic range but could also return data that is located only 5’ of the queried start. Ensuring the end of a returned record is greater than or equal to the queried start further restricts the query. The range query performs an order of magnitude faster than querying with just a start and end and performs well in data volumes ranging from 1bp to 1,000,000bp.

**Figure 3:**
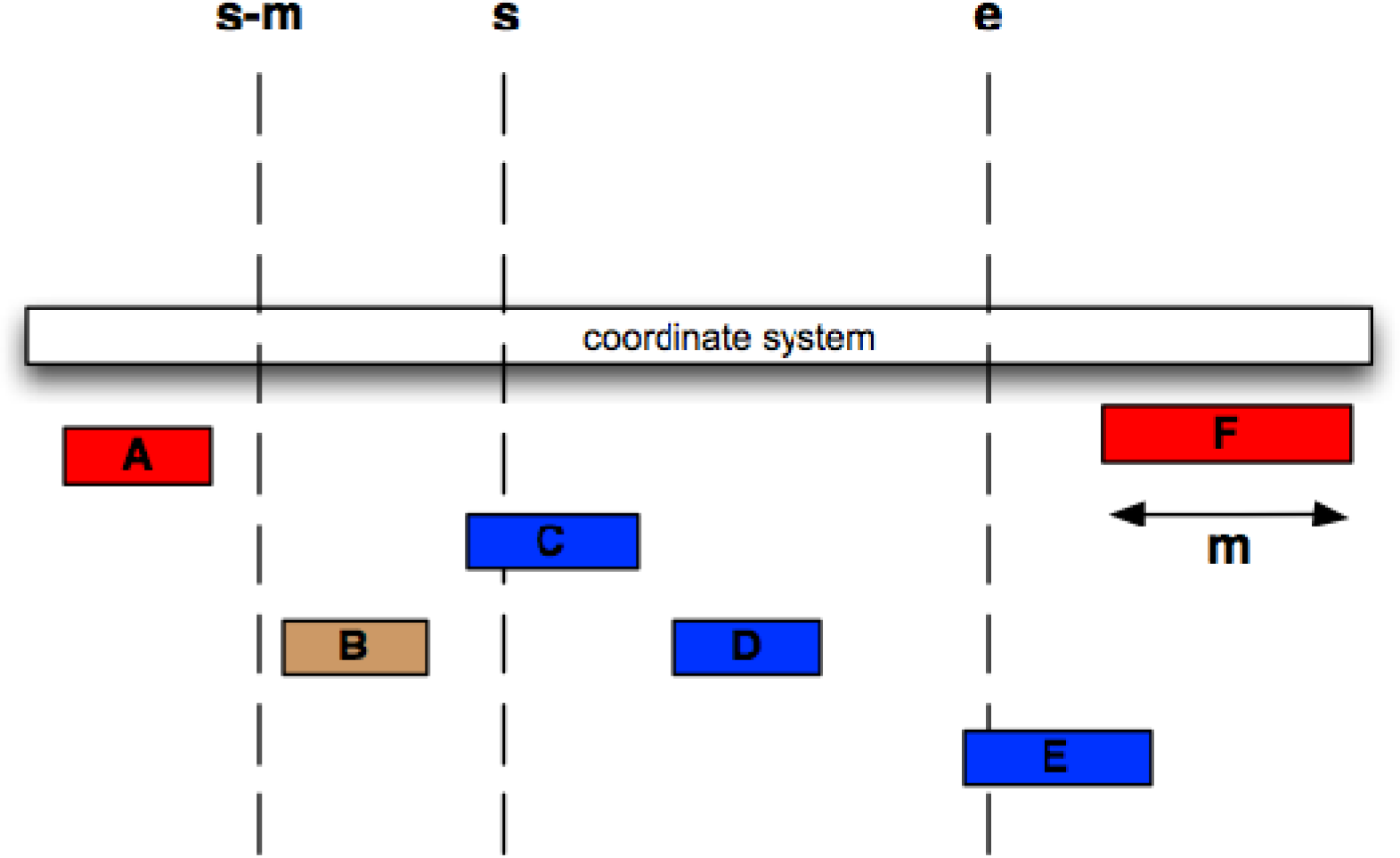
Efficient Searching of genomic features. We wish to find all features between coordinates *s* and *e* (i.e. C, D and E). The maximum length of a feature for this coordinate system is m, i.e. the length of feature F. We therefore extract all features whose start lies between s – m and e, then exclude B, since it ends before s.

### Ensembl Database Registry

Before retrieving data from the Perl API it is essential to locate all databases to be queried. Depending on the API call this could be a single or multiple databases. The Ensembl database registry solves this problem by providing two major functions. First, it is a global associative array that holds a reference to data adaptors keyed by species name, database type and data type. The registry also holds a list of species name aliases allowing the use of common terms such as “human” to find a *Homo sapiens* adaptor. Second, the registry can automatically populate itself from a MySQL server (requiring a hostname and username be specified) and it searches for databases corresponding to a single release. Additional routines enable searching across multiple MySQL instances or from an external configuration file.

### Alternative Data Stores

In recent years, projects such as ENCODE (26), 1000 Genomes (10) and Illumina BodyMap 2.0 have produced a large number of genomic data sets. This situation has given rise to large indexed file formats including BigBed, BigWig (27) and BAM (25). We now use these data stores when existing database solutions fail to scale adequately. The data contained in these files is normally quite static between releases and not relational.

We have developed a metadata file storage system for locating these files called DataFiles. A data file entry describes the assembly version a file maps to, the analysis method used to generate the data, a name, the file type and an optional versioning flag. The DataFiles API can construct a POSIX filename path from this information and provide an object capable of reading this data. For example, BAM files trigger creation of an object with bindings to HTSlib (http://www.htslib.org). DataFiles currently supports BAM, BigWig, BigBED and a specialised format BAMCOV, which combines a BAM file and pre-computed read coverage held in a BigWig file.

## Data management for non-vertebrate genomes

### Multi-species databases

The Ensembl Core infrastructure was designed for vertebrate genome assemblies in general and human in particular and included an assumption that each species would be contained within a single database. As the number of genomes grew, putting each in its own database was putting significant pressure on our MySQL infrastructure and may have eventually exposed underlying file system limitations. To avoid these problems, we devised a method to store multiple genomes in a single Core database. For example, Ensembl Bacteria release 32 (August 2016) contains 41,610 species of bacteria and archaea held in only 173 databases.

Multi-species support involves associating separate coordinate systems to each species. Since every genomic annotation in an Ensembl database is mapped on a sequence region that is itself linked to a coordinate system, selecting the appropriate coordinate system will create species-specific queries. The Core API transparently handles the storage and querying of these multi-species databases by joining back to *seq_region* and *coord_system* tables when required. The API avoids these extra steps when dealing with a single-species database, but on a multi-species database (characterised by a name that includes the term ‘collection’) it runs the required queries to discover and register the species contained. This naming convention is not required if manually working with a multi-species database. Controlled vocabulary tables are considered global to all species within a single database. Metadata linked to a species, such as its name, are held in the *meta* table and are linked to its species identifier.

### Circular chromosomes

Prokaryote species require native support for circular genomes. Specifically, on sequence regions marked as circular, a feature that passes through the origin of replication (ori) will have an end coordinate “lower” than its start coordinate. We require that a feature can pass through the ori only once. Ori spanning queries are decomposed by the Core API into two queries: the first from the requested start coordinate to the sequence region’s length and the second from position 1 to the requested end coordinate. Features appearing in results of both queries (i.e. those that pass through the ori) are returned once.

### Polycistronic Transcripts

Prokaryote and some eukaryote species also require native support for polycistronic genes. These are genes that are transcribed as a single polycistronic mRNA and undergo trans-splicing to result in multiple polypeptide products (28, 29). The Ensembl database schema was extended to represent polycistronic transcripts using three tables. The *operon* and *operon_transcript* tables locate the full range of polycistronic transcripts available and group them under a single regulatory element. Each operon_transcript record is linked to a set of polypeptide encoding genes via the *operon_transcript_gene* table thereby annotating the available proteins. Every polypeptide encoded by a polycistronic transcript is held as a record in the gene and transcript tables. This allows the Ensembl Core API to translate the resulting transcript into a protein sequence without significant re-engineering. Polycistronic data is currently available for *Escherichia coli str. K-12 substr. MG1655*.

## Projecting data between assembly versions and gene set updates

Periodically new assemblies are released for species that have previously been annotated in Ensembl. As these normally incorporate new sequence data and are generally of better quality than previous assembly versions, they tend to be quickly adopted for new genomic analysis efforts. However, because a new assembly will have a different coordinate system, directly comparing the results of analyses on two assembly versions is difficult or impossible. The Ensembl Core database and API store and manage assembly and identifier mapping data to help transition and compare results between assemblies.

### Mapping between assemblies

We first generate a mapping from the old to the new assembly that takes advantage of shared contigs between the two assembly versions. The contig order, within each chromosome, is compared between versions. If contig accessions and versions are maintained between assembly versions, we look at the versioned INSDC accession; otherwise we generate a MD5 checksum of the underlying sequence. Neighbouring contigs with conserved order and orientation are collapsed into a single mapping held at the chromosome level. Gaps in the mapping between chromosomes are filled in by a LASTZ alignment (30). Finally, the resulting mappings are recorded in the assembly table.

Once a mapping is available between two assemblies, annotations can be transferred from one to another taking into account any insertions or deletions of DNA sequence. This is available from our Perl API via two subroutines: *project* to report a target coordinate set and *transform*, which generates a new annotation object in the target coordinates. Both are available on any Ensembl Feature class including genes, transcripts, repeats and variants. These subroutines are used when annotating an updated assembly to transfer the existing annotation onto the new assembly for quality control (7). As such, our method attempts to minimise gaps between assemblies and help map gene annotation cleanly between assemblies.

In addition we create chain format files of our assembly version mappings to be used by liftover (31) or CrossMap (32). The latter is utilised for our online Assembly Mapper tool (33). Chain files are available from our FTP site and include additional mappings that use the UCSC naming scheme for regions (i.e. chr1 rather than 1).

### Mapping stable identifiers

As new experimental sequence evidence becomes available or when a new genome assembly for a species is published, it is possible to generate updated and more accurate gene annotation. In these cases, linking new gene models to their older versions is particularly useful for reproducibility and other analyses.

The Ensembl gene annotation system uses stable identifiers to uniquely identify each feature across releases. The stable ID mapping pipeline ensures the identity of these features is kept consistent across releases, whereas the version is incremented each time a feature is modified. For example, if a transcript in Ensembl release 84 has a longer UTR region than the same transcript in release 83, the version will be different but the stable id will be the same. As the Ensembl gene annotation is associated directly with an assembly, our challenge is to identify the gene and transcript models that are found on both assemblies and transfer our identifiers between those models. Other Ensembl resources such as GeneTrees and Ensembl Protein Families also use stable identifiers with procedures adapted to the specific data types (3).

Our methodology is based on hierarchical identity, starting from exon overlaps: all exons from the source set are compared to all the exons from the target set. The comparison is based on coordinate overlap when both source and target are on the same assembly, and based on sequence alignment when going from one assembly to another. If a pair of exons share 90% or more of their bases they are considered the same and the exon stable id is transferred from source to target. Thus, when using coordinate overlap on the same assembly the new exon must share 90% of the bases to retain the stable id and when comparing between assemblies we require 90% identity in the alignment. The exon stable ID version is incremented if the coordinates or the sequence is not a perfect match.

Transcript identifiers are then mapped by comparing sets of exons (as defined by the source and target annotation) and looking for sets with 25% or greater intersection. Matches are penalised when the compared transcript functions do not match each other to reduce the chances of incorrect transfer of identifiers. A simple change in function will incur a 10% penalty increasing to 20% if that change causes a significant alteration in assigned function. For example, changing a transcript’s function from short nuclear RNA to short cytoplasmic RNA will only incur a 10% penalty since both of these functions are linked to small non-coding RNAs. If a transcript changes from coding to non-coding we apply a 20% penalty since this represents a significant change in assigned function. A transcript version will increment when there is a change in the underlying cDNA sequence or exon-splicing pattern. The identity of a transcript is thus defined by the list of its exon coordinates and its underlying sequence.

Protein identifiers are mapped via the underlying transcript mapping and versions are incremented when protein sequence changes.

Genes identifiers are mapped based on the shared transcripts between source and target data sets. Any stable identifier or version change in the underlying transcripts will cause an increment in the gene’s stable identifier version.

In some cases, a source feature can map with comparable scores to two or more features. These ambiguous cases are resolved by comparing the biotypes, the parent and children features or the internal ids. If it is not possible to clearly identify the best candidate, all existing IDs are discarded and new stable IDs are assigned (Figure 4).

**Figure 4:**
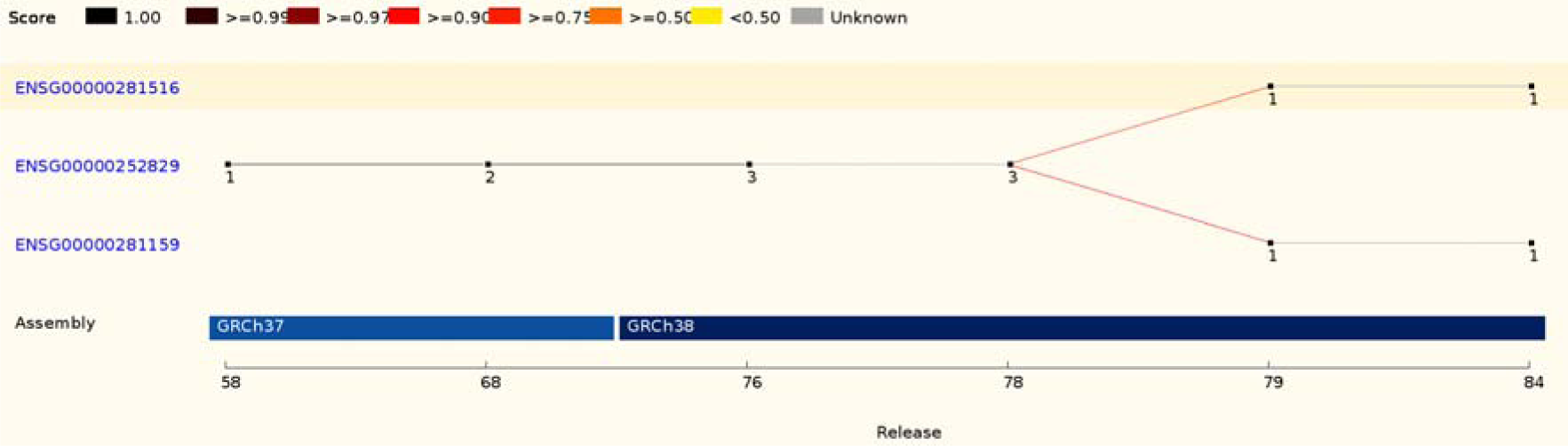
This ID History Map for the SCARN4 gene (http://www.ensembl.org/Homo_sapiens/Gene/Idhistory?g=ENSG00000281516) aligns Ensembl release numbers, genomic assembly versions, and version numbers of that gene across multiple Ensembl IDs. The different updates in the version ID are represented as a chain of small nodes, connected by lines. The colour of the line reflects how well consecutive versions match, for recent releases. If a score was not calculated (typically in older releases), the line is grey.

## External references

Linking Ensembl entities such as genes, transcripts and proteins to external resources such as UniProt, RefSeq, RNACentral (34) and HGNC increases the value of our annotation by enabling cross resource querying.

We employ four methods to perform this linkage: direct (mappings provided by a third party), location overlap (annotation overlapping the same genomic location), checksum matches (MD5 digests of sequences) and alignment executed by exonerate (35). Alignment based linkages can be performed between any resource where sequence is attached to an identifier. After our sequence sets have been aligned against an external set, results are limited to the best hit or a range of hits such as the top five. Hits are filtered by their percentage identity and all alignments are subject to a minimum threshold of identity. If we manage to align multiple sequences to the same degree of identity then all are linked. A selection of the resources linked and the mapping method is shown in Table 1.

**Table 1:** A selection of resources mapped by the Ensembl cross reference system indicating the methods used to map, the Ensembl feature mapped to, any additional resources brought in by this association and whether the resource is used to name genes.

| Resource | Mapping Method | Linked To | Transitive Resources | Used for naming |
| --- | --- | --- | --- | --- |
| HGNC | Direct | Genes | - | Yes |
| UniProt | Direct, alignment | Proteins | PDBe | Yes |
| RefSeq mRNA | Location overlap, alignment | Transcripts | EntrezGene | Yes |
| RefSeq Proteins | Alignment | Proteins | - | No |
| UniProt Archive (UniParc) | Checksum | Proteins | - | No |
| RNACentral | Checksum | Non-coding transcripts | - | No |
| MGI | Direct | Genes | - | Yes |

Our approach selects the best matches according to a priority rating. Direct links are assumed to have the highest priority followed by location, checksum and then alignment based links. Transitive links are also imported including links from UniProt to PDBe (36). Once all links are assembled we use these to select a name for each gene. If a species has a nomenclature committee such as HGNC or MGI (37) these links will take the highest priority. Otherwise we revert to using RFam (38), mirbase (39), UniProt or EntrezGene (40) names and, if there is no other option, a clone name. Links are then stored with gene, transcript and protein records. If a link was generated by alignment we retain the percentage identities of the mapping and a CIGAR string representation of the alignment. Alternatively if the link was from an ontology, we record the three letter evidence code assigned to the link as described by the Gene Ontology Consortium such as IEA (Inferred from Electronic Annotation) for unreviewed annotation assigned by a computational method (41).

## Discussion

Over the past several years, we have engineered substantial enhancements to our API without significantly changing the way existing developers interact with it via existing public interfaces. Updates such as multi-species and polycistronic transcript support necessitated a number of schema and code changes, but created minimal API changes. For example, circular genome support required API changes only, while polycistronic transcript support required changes to the database schema but did not alter existing API methods. None of these updates required existing code to be modified, except to use a new feature. Indeed, Ensembl’s gene orthology and paralogy annotation pipeline (42) was run on multi-species core databases with no modifications. To implement these changes with such a minimal level of impact requires the level of abstraction described above.

Schema changes are made only when necessary and incompatible changes are applied when we feel the current schema falls short of its intended goals. Where possible, migration plans are offered between schemas for a number of releases, giving developers time to migrate. Such was the solution offered when stable identifier tables merged with their parent tables in Ensembl release 65. Where a method is considered no longer applicable or supplanted by a superior method the original is deprecated and developers are pointed to a replacement. An extensive set of unit tests (4,646 in Ensembl release 84) ensures that code development does not cause a break in our expected API behaviour. Due to the API’s ubiquitous usage within our data production cycles, performance regressions are generally quickly identified and corrected.

The API continues to support novel methods of information delivery across multiple species as highlighted by our REST Service (43). By building on the Perl API, the REST API has a solid platform for development and a stable robust model of data transfer. It is compatible with any Ensembl infrastructure based resource including Ensembl Genomes (44), which has released its own installation of this service. We see the exposure of Ensembl data to alternative programming languages as an important development.

The flexibility of the Ensembl Core infrastructure is most clearly displayed by the many large-scale projects that reuse it, including Ensembl Genomes, Gramene (45), WormBase and WormBase ParaSite (46), VectorBase (47), PomBase (48) and AvianBase (49). The versatility of the Ensembl Core software infrastructure, including the Perl and REST APIs, is further demonstrated by the third party tools which incorporate and extend it (50–52). These vibrant and active research communities regularly bring in new demands and requirements that, together with the challenges of new data types described above, ensure the development of our Core infrastructure is as active now as it has ever been.

## Conclusions

The Ensembl core API and database is a highly extensible, performant, adaptable and consistent interface to genomic data. This has allowed other projects to create their own resources using the Ensembl infrastructure. The Perl APIs continue to be the primary mechanism of data access within the project, supporting the Ensembl Website and our newer data distribution mechanisms such as the Ensembl REST API. As more species are sequenced and the amount of species-specific data increases, the foundation of the Ensembl core infrastructure will ensure successful scaling to meet the future needs of the genomics community.

## Acknowledgments and funding

We acknowledge the contributions of former team member and original Ensembl Core Software development leader Arne Stabenau and from the rest of the Ensembl team. We thank Guy Coates, Peter Clapham and Tim Cutts for maintaining the Ensembl computer systems.

This work was supported by the Wellcome Trust (grant number WT108749/Z/15/Z), which provides majority funding to Ensembl; the Biotechnology and Biological Sciences Research Council (BB/L024225/1, BB/I025506/1 and BB/I025360/2), and the European Molecular Biology Laboratory. Funding for open access charge: The Wellcome Trust.

